# Abnormal molecular signatures of inflammation, energy metabolism and vesicle biology in human Huntington disease peripheral tissues

**DOI:** 10.1101/2022.02.03.475821

**Authors:** Andreas Neueder, Kerstin Kojer, Tanja Hering, Daniel J. Lavery, Jian Chen, Nathalie Birth, Jaqueline Hallitsch, Sonja Trautmann, Jennifer Parker, Michael Flower, Huma Sethi, Salman Haider, Jong-Min Lee, Sarah J. Tabrizi, Michael Orth

## Abstract

**Background:** A major challenge in neurodegenerative diseases concerns identifying biological disease signatures that track with disease progression or respond to an intervention. Several clinical trials in Huntington disease (HD), an inherited, progressive neurodegenerative disease, are currently ongoing. Therefore, we examined whether peripheral tissues can serve as a source of readily accessible biological signatures at the RNA and protein level in HD patients.

**Results:** We generated large, high-quality human datasets from skeletal muscle, skin and adipose tissue to probe molecular changes in human premanifest and early manifest HD patients – those most likely involved in clinical trials. In-depth single nucleotide polymorphism data across the *HTT* gene will facilitate the use of the generated primary- and iPSC cell lines in allele-specific targeting approaches. The analysis of the transcriptomics and proteomics data shows robust, stage-dependent dysregulation. Gene ontology analysis confirmed the involvement of inflammation and energy metabolism in peripheral HD pathogenesis. Furthermore, we observed changes in the homeostasis of extracellular vesicles, where we found consistent changes of genes and proteins involved in this process.

**Conclusions:** Our ‘omics data document the involvement of inflammation, energy metabolism and extracellular vesicle homeostasis. This demonstrates the potential to identify biological signatures from peripheral tissues in HD suitable as biomarkers in clinical trials. Together with the primary cell lines established from peripheral tissues and a large panel of iPSC lines that can serve as human models of HD, the generated data are a valuable and unique resource to advance the current understanding of molecular mechanisms driving HD pathogenesis.

## Introduction

The CAG repeat expansion mutation in exon 1 of the huntingtin gene (*HTT*) causes Huntington disease (HD), a progressive movement disorder with dementia and behavioral abnormalities [1]. Huntingtin is expressed ubiquitously throughout the body, and HD does not exclusively affect the brain. In addition to a particular vulnerability of striatal and cortical neurons to the mutation, several lines of evidence suggest that mutant HTT influences molecular function in peripheral tissues including skeletal muscle, peripheral blood monocytes, liver and others [2]. Hence, the investigation of peripheral tissues and cells holds promise to reveal important insight into the physiological function of HTT that may include transcriptional processes, protein trafficking and vesicle transport [3, 4]. It can also reveal peripheral signatures of HD that track with the evolution of the expression of HD in the central nervous system. This could have potential as an easily accessible and low invasive biomarker in clinical trials in HD. Finally, primary cell cultures established from peripheral tissues such as skin can be used to generate pluripotent stem cells, which, when differentiated into neuronal and non-neuronal cells can serve as human models of HD.

The objective of the Multiple Tissue Monitoring in Huntington disease (MTM-HD) study was to comprehensively examine the molecular biology at the RNA and protein level of peripheral tissues from *HTT* CAG repeat expansion mutation carriers and sex and age matched healthy volunteers. All participants were also participants in Enroll-HD (www.enroll-hd.org), and extensive demographic and phenotypic data were collected according to the Enroll-HD protocol. The analyses of proteomics and transcriptomics datasets in conjunction with the available deep clinical phenotype information can give valuable insights into more complex signatures in human carriers of the HD gene mutation and provide a resource of peripheral tissues and primary cells with accompanying multi-layer ‘omics datasets.

The main potential of these types of analyses is in relating biological signature(s) – either in individual modalities and/or tissues or across modalities and tissues – with CAG repeat length as the severity of the expression of the HD gene mutation. Lastly, the insight gained through these analyses can inform future research with the aim to better understand the nature of any peripheral biological signature and the impact an intervention, e.g. in a clinical trial, can have on them.

## Results

Skeletal muscle, adipose tissue and skin from an open biopsy of the *quadriceps femoris* muscle, as well as blood, were collected from 20 healthy controls, 21 pre-symptomatic and 20 early motor manifest HD (UHDRS total functional capacity stages 1 and 2) patients. Additionally, primary fibroblast and myoblast cell lines have been established. In the following we present the demographic and clinical data for the participants, the initial findings of transcriptional and proteomic dysregulation in tissues, as well as the analysis of the generated fibroblasts by RNAseq and *HTT*-centered single nucleotide polymorphism (SNP) sequencing. The analysis of the epigenetic clock in the fibroblast lines has already been published [5]. As an additional resource, pluripotent stem cell lines (iPSCs) have been generated from the primary fibroblast lines and are available for further studies.

### Participant’s demographic and clinical data

At both sites, a total of 24 unaffected individuals (control), 23 pre-manifest (known mutation carriers, before disease onset; pre-HD) and 21 early HD (early-HD) patients were recruited into the study that was approved by the Ethics Review Boards of Ulm University and University College London. Following a detailed description of the study participants gave informed consent according to the Declaration of Helsinki. Participants attended two visits: the screening and sampling visits, which were no more than 1 month apart. At the screening visit, participants were assessed for inclusion and exclusion criteria (Table 1). Ultimately, 20 control, 21 pre-HD and 20 early-HD participants were included in the sampling visit of the study. Summary data for these 61 participants are shown in Table 2. For the full dataset including statistics see supplementary file (MTM_all_metadata.xlsx). Participants from all three groups were evenly recruited across the two sites, and were matched for gender, as well as age (Table 2). Neither age, weight, height, nor BMI were significantly different between the groups (one-way ANOVA). CAG repeat sizes of the expanded allele were not significantly different in the pre-HD and early-HD groups while DBS was higher in the early-HD group than in the pre-HD group, as expected (Table 2). All clinical measures were only significantly different in the early-HD group compared to the control or pre-HD group (Table 2 TFC, TMS, FA, IS; one-way ANOVA with Bonferroni *post hoc* test, *p* < 0.001).

**Table 1.** Inclusion and exclusion criteria for participants of the MTM-HD study.

| Inclusion criteria | Exclusion criteria |
| --- | --- |
| Age 30-50 | Major psychiatric disorder at time of enrolment |
| Caucasian ethnicity to minimize genetic differences in this relatively small sample set | On long-term anti-coagulants e.g. Warfarin for mechanical heart valves, pulmonary embolism or atrial fibrillation |
| Huntington disease gene CAG repeat length 40-55, for gene expanded subjects only | On combination of Aspirin and Clopidogrel for ischaemic heart disease |
| Premanifest subjects stratified as closer to predicted disease onset (disease burden score >250, UHDRS diagnostic confidence score <3) | Any relevant condition (including presence of psychosis and/or confusional states), behavior, laboratory value or concomitant medication which, in the opinion of the investigator, makes the subject unsuitable for entry into the study |
| Early HD subjects (Stage 1 & 2 disease as quantified by UHDRS total functional capacity score 7-13) aged 30-50 | Recent major abdominal or limb surgery within last month |
| Healthy volunteers aged 30-50 | Fear of needles |
| REGISTRY or ENROLL-HD assessment within 6 months of screening visit, for gene expanded subjects only | Anaemia (Hb<11.5), thrombocytopenia (platelets <150) or coagulopathy (INR >1.5) identified on screening and/or blood tests |
| Presence of Companion to accompany them home | High-dose nutraceuticals e.g. creatinine, co-enzyme Q, vitamin E |
| Capable of providing informed consent | Significant Lower limb impairment of any kind |
| Subjects must have no clinically significant and relevant history that could affect the conduct of the study and evaluation of the data, as ascertained by the investigator through detailed medical history | Significant cardio-respiratory or other medical comorbidities such as Ischaemic Heart Disease, Diabetes, Chronic Obstructive Pulmonary Disease, Chronic Liver Disease, Stroke with hemiparesis, BMI >30 and history of malignancy in last two years |
| Ability to tolerate procedures, fasting and blood sample donation | Illicit drug use and/or alcohol abuse > 21 units/wk males; 14 units/wk female |
|  | Participation in an investigational drug trial within 30 days of screening visit |
|  | Pregnancy |
|  | Allergy/previous adverse reaction to local anesthetic agents |

**Table 2.** Summary data for the 61 participants of the MTM-HD study.

| HD status | Site | Gender | Age [years] | Short CAG allele | Long CAG allele | DBS | Weight [kg] | Height [cm] | BMI | TFC | TMS | FA | IS |
| --- | --- | --- | --- | --- | --- | --- | --- | --- | --- | --- | --- | --- | --- |
| control | U: 10<br>L: 10 | f: 10<br>m: 10 | 41.3<br>[31.6;<br>55.9] | 15.2<br>[10;20] | 18.5<br>[15;27] | n. a. | 74.4<br>[45.3;102] | 171.7<br>[158;187] | 25.1<br>[17.6;33.7] | 13<br>[13;13] | 0.5<br>[0;4] | 25<br>[25;25] | 100<br>[100;100] |
| pre-HD | U: 11<br>L: 10 | f: 10<br>m: 11 | 41.6<br>[31.2;<br>56.5] | 19.6<br>[9;24] | 43.6<br>[40;48] | 327.7<br>[158.6;<br>423.8] | 76.2<br>[49.2;115] | 172.8<br>[154;190] | 25.4<br>[20.4;35.5] | 12.7<br>[11;13] | 5.3<br>[0;18] | 24.7<br>[22;25] | 97.6<br>[80;100] |
| early-HD | U: 10<br>L: 10 | f: 10<br>m: 10 | 45.4<br>[35.4;<br>58.9] | 18.9<br>[9;25] | 44.5<br>[41;50] | 406.7<br>[281.0;<br>657.3] | 73.7<br>[55;94.4] | 173.5<br>[153;197] | 24.4<br>[18.6;27.9] | 10.9<br>[7;13] | 24<br>[8;49] | 22.3<br>[16;25] | 88.4<br>[70;100] |

Data are mean and minimal and maximal values, respectively, in brackets. Abbreviations: Site: U = Ulm; L: London. Gender: f = female; m = male. DBS = disease burden score (not applicable (n.a.) in the control group). BMI = body mass index. TFC = total functional capacity. TMS = total motor score. FA = function analysis. IS = independence scale.

### *HTT* gene SNP sequencing

An important consideration in therapeutic *HTT* lowering trials is preservation of a certain level of HTT protein. While inactivation of HTT in the brain of adult mice [6] and primates [7] seems to be tolerated for some time, inactivation of HTT beyond a certain level, likely 50% of the normal level, in the developing, as well as the adult brain potentially has severe consequences (reviewed in [8, 9]). Therefore, an attractive approach is to target the alleles of single nucleotide polymorphisms (SNPs) that are phased on the mutated *HTT* allele. To facilitate such approaches, we genome-sequenced the primary fibroblast lines generated in this study by the 10X Genomics Chromium technology. We only sequenced the lines that were generated from HD patients (n = 39). Due to patient privacy and legal reasons, here we only show the *HTT* gene (± 100 kbp) associated SNPs (datafile in the supplementary information; MTM-HD_HTT_SNP_haplotyping_fibroblasts.xlsx). Overall, 98.3% of all identified SNPs could be assigned to a specific haplotype. The largest block of assembled *HTT* sequencing data was around 14.5 million base pairs.

We previously defined 16 haplotypes that are common in HD subjects with European ancestry [10], and demonstrated allele-specific target sites for each haplotype [11, 12]. To maximize the discrimination power of the haplotypes and to cover non-Europeans, we extended the *HTT* haplotype definitions by using 1000 Genomes Project (KGP) data (phase 3, all populations). Based on the same 21 genetic variations that were used to define the original 16 haplotypes [10], 253 additional haplotypes were identified and subsequently assigned based on the allele frequency in all KGP populations. For example, excluding the original 16 haplotypes, hap.17 represents the most frequent *HTT* haplotype in KGP data, accounting for approximately 3% of haplotypes in KGP chromosomes. The list of all newly defined HD haplotypes can be found in the supplementary information (MTM-HD_HD_haplotype_definitions.xlsx).

Out of the total of 39 sequenced samples, the *HTT* haplotypes of 37 samples could be conclusively defined (Fig. 1A and B). A phylogenetic tree analysis on the *HTT* haplotypes to study evolutionary relationships between the different *HTT* alleles revealed three major groups of *HTT* haplotypes (represented by colors in Fig. 1C and D).

**Fig. 1.**
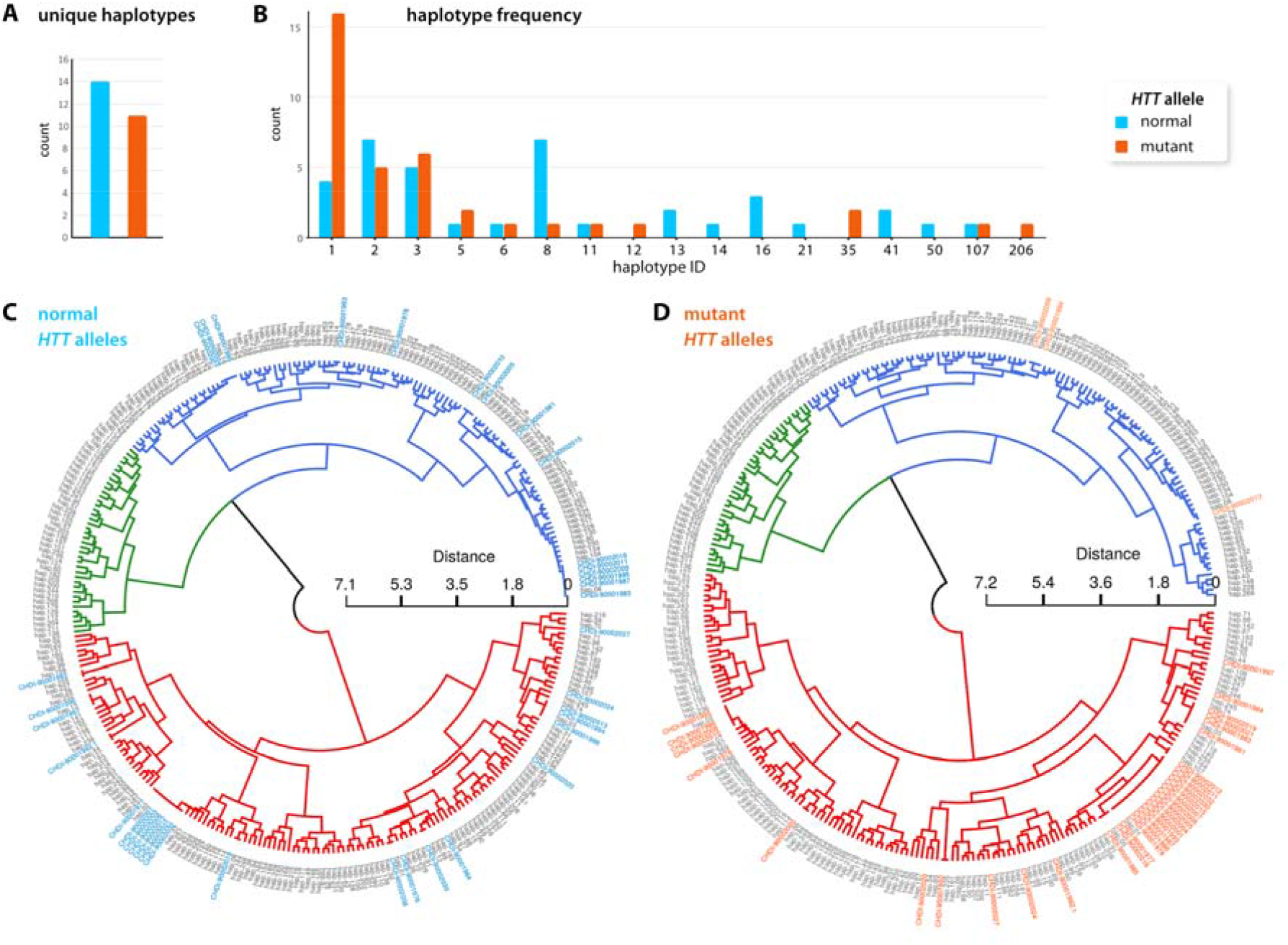
HD haplotype analysis. (**A**) Uniquely detected haplotype count for the normal (light blue) and the mutant (orange) allele in the sequenced patient samples. (**B**) Histogram plots of HD haplotypes distribution of normal and mutant *HTT* alleles. Exact and nearest neighbor matches were considered. A phylogenetic dendrogram analysis of the HD haplotypes of normal (**C**) and mutant (**D**) *HTT* alleles highlights how closely the SNPs are related. The MTM-HD samples were assigned a ‘virtual’ haplotype and added to the analysis. Therefore, MTM-HD samples cluster together with their most closely related haplotype, which also corresponds to the assigned haplotype as shown in **A**. The HD haplotypes are indicated as hap.X and the MTM-HD fibroblast samples are shown by their CHDI ID. Colors of the dendrograms represent the three main haplotype groups as defined by the k-mer clustering. MTM-HD samples were colored according to their expansion status. See also supplementary table for full data.

Mutant *HTT* alleles fell into only 2 of these major groups (blue and red). Notably, the haplotypes in the blue haplotype group seemed to be less prone to harbor an expanded *HTT* allele (compare the frequency of HD patient samples (CHDI ID) in the blue clade in Fig. 1C and D): 15 of the normal *HTT* alleles and only 3 of the mutant alleles were attributed to the blue haplotype group, while 22 of the normal *HTT* alleles and 34 of the mutant alleles were attributed to the red haplotype group (Fig. 1C and D).

### Generation of iPSC lines

To generate a flexible model for HD a total of 45 iPSC lines were derived from the primary fibroblast cell lines in this study (controls and HD patients). The iPSC lines are available from the European Bank for Induced pluripotent Stem Cells (EBiSC; search for ‘CHDI’ at https://cells.ebisc.org). Re-programming of fibroblasts was conducted using the non-integrative Sendai virus strategy expressing KLF4, MYC, POU5F1 (OCT4) and SOX2 [13]. All lines were characterized regarding expression of pluripotency markers (OCT-4 (POU5F1), TRA-1-60, SSEA-1 and SSEA-4), retention of re-programming vector sequences, morphology and karyotype. Furthermore, iPSC lines were differentiated into endo-, meso- and ectodermal cells and marker expression was analyzed (endoderm: CXCR4, GATA6, SOX17; mesoderm: MIXL1, NCAM1, VIMENTIN; ectoderm: HES5, NEUROD1, PAX6). All data are freely available from the EBiSC website (at the time of June 2021).

### Transcriptomic analysis of adipose and muscle tissue

We next analyzed the transcriptome of adipose and muscle tissue from the biopsies. We generated 50 bp long, paired-end reads with a depth of approximately 140 to 200 million reads per sample of rRNA depleted total RNA (See also Method section for more detail and supplementary information for the RNAseq count files; MTM-HD_RNAseq_TISSUE_outliers_removed_normalized_counts.xlsx). In total we found 78 genes in adipose (Fig. 2A) and 53 genes (Fig. 2B) in muscle to be significantly dysregulated (Benjamini-Hochberg adjusted *p*-value < 0.05). We only considered genes with counts in at least 9 samples in at least one group (50%), in at least one comparison (pre-HD vs. controls; early-HD vs. controls; early-HD vs. pre-HD). The full analysis files can be found in the supplementary information (MTM-HD_RNAseq_TISSUE_outliers_removed_DESeq2_analysis.xlsx).

**Fig. 2.**
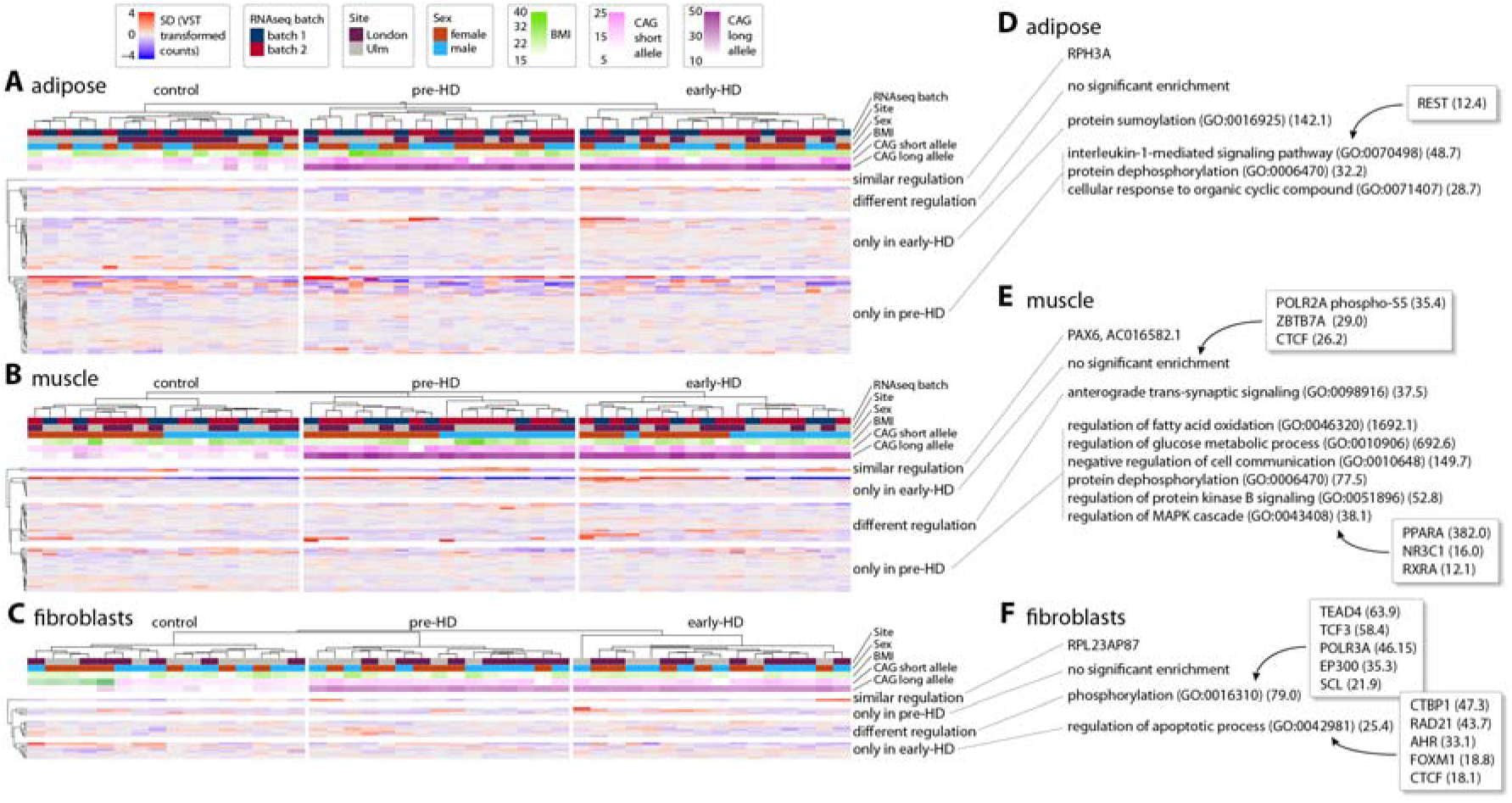
Transcriptomics analysis of adipose and muscle tissue, and fibroblast lines. (**A-C**) The deviation of variance stabilized counts (VST) from the mean of all samples are shown for each sample (columns). Genes are clustered according to their regulation as defined by the Benjamini-Hochberg adjusted *p*-value (< 0.05) in the DESeq2 analysis. Only genes with counts in 9 or more samples in at least one group for at least one comparison (pre-HD vs. controls; early-HD vs. controls; early HD vs. pre-HD) are shown. Co-variates (RNAseq batch, site of sampling (Site), Gender, BMI and the CAG allele sizes) are shown above the expression matrix. (**A**) 78 genes in adipose were significantly dysregulated. (**B**) 53 genes in muscle were significantly dysregulated. (**C**) 21 genes in the fibroblast lines were significantly dysregulated. (**D-F**) Gene ontology enrichment analysis with Enrichr (see methods). Non-redundant biological process enrichments (GO Biological Process 2018) and potential regulators (ChEA 2016, ENCODE 2015, TTRUST 2019) are shown. Only terms with *p* < 0.05 (Fisher exact test) and at least 2 genes for the enrichment were considered. The combined score of enrichment is shown in brackets. GO enrichment for adipose (**D**), muscle (**E**) and fibroblast (**F**) RNAseq data. See also supplementary tables for full data.

Notably, only very few genes were regulated in the same way in the pre-HD and early-HD groups when compared to controls (Fig. 2A and B, similar regulation). Several genes were differentially regulated in both groups (Fig. 2A and B, different regulation). One gene in this group, *TBC1D3D* (ENSG00000274419), was found to be differentially regulated in both tissues. In both tissues, the gene was upregulated in the early-HD group compared with the pre-HD group (adipose: log2-fold change 4.47, p_adj_ = 0.003; muscle: log2-fold change 1.51, p_adj_ = 0.041). In both tissues sets of genes were exclusively dysregulated in either the pre-HD or early-HD groups (Fig. 2A and B, pre-HD only and early-HD only).

Gene ontology enrichment in adipose tissue for the different comparisons showed that REST (RE1-Silencing Transcription factor) target genes are dysregulated only in the pre-HD group (Fig. 2A), while protein sumoylation was only dysregulated in the early-HD group (Fig. 2A). Alteration in protein sumoylation is a well described mechanism in HD, both for HTT itself, as well as in general [14-18]. Also, REST has been implicated in HD pathogenesis [19-21].

Gene ontology enrichment analysis in muscle points towards disrupted homeostatic pathways, especially in the pre-HD group (Fig. 2B). Interestingly, *PAX6*, an important regulator of diverse peripheral and central nervous system processes, was highly and progressively up-regulated in both HD groups (Fig. 2B, similar regulation), suggesting a compensatory mechanism of muscle regeneration in response to mutant HTT expression. Similar to our finding, in the muscle of R6/2 mice, a higher level of satellite cells has been found, concomitant with an increase in PAX7, another member of the PAX transcription factor family [22]. The enrichment analysis of the dysregulated genes in the muscle pre-HD group suggested PPARA (peroxisome proliferator activated receptor alpha) as a regulatory protein with very high confidence (Fig. 2B).

### Transcriptomic analysis of fibroblast lines

We generated 100 bp long, paired-end reads with a depth of approximately 40 to 60 million reads per sample of polyA-enriched total RNA (For RNAseq count files see supplementary information; MTM-HD_RNAseq_fibroblasts_outliers_removed_normalized_counts.xlsx). Evaluation was performed in the same way as described for tissues above. Only few genes were significantly dysregulated between pre-HD and early-HD groups as compared to controls (Fig. 2C). Consequently, gene ontology analysis resulted in few enriched terms (Fig. 2C). Notably, *TBC1D3D* (ENSG00000274419), which was found to be dysregulated in both, the adipose and muscle datasets (see above and Fig. 2A and B), was also dysregulated in the fibroblast dataset. Its expression pattern was consistent between all three datasets with a down-regulation in the pre-HD group and a restoration towards control levels in the early-HD group.

### Proteomic analysis of skin, adipose and muscle tissue

Next, we generated proteomics data from the skin, adipose tissue and muscle biopsies. In adipose tissue, we identified 1347 proteins, in muscle 2671 proteins and in skin 4640 proteins. After correction for confounding variables (TMT-plex, site of sampling, gender, age and BMI; see Methods section for details), we performed dysregulation analysis using ROTS (reproducibility-optimized test statistic) [23]. ROTS uses a non-predetermined optimized test statistic for each dataset by utilizing bootstrapped datasets that preserve the individual top-ranked features and maximizing the overlap between those. The data files for peptide and protein analysis can be found in the supplementary information (MTM-HD_proteomics_TISSUE_peptide/protein.xlsx). The analyses are shown as volcano plots for each of the pairwise comparisons for each tissue in Fig. 3 with significantly dysregulated proteins (*p*-value < 0.001) highlighted. Proteins that were also determined to be dysregulated based on analysis of permutated sample lists are highlighted in bold (FDR < 0.05).

**Fig. 3.**
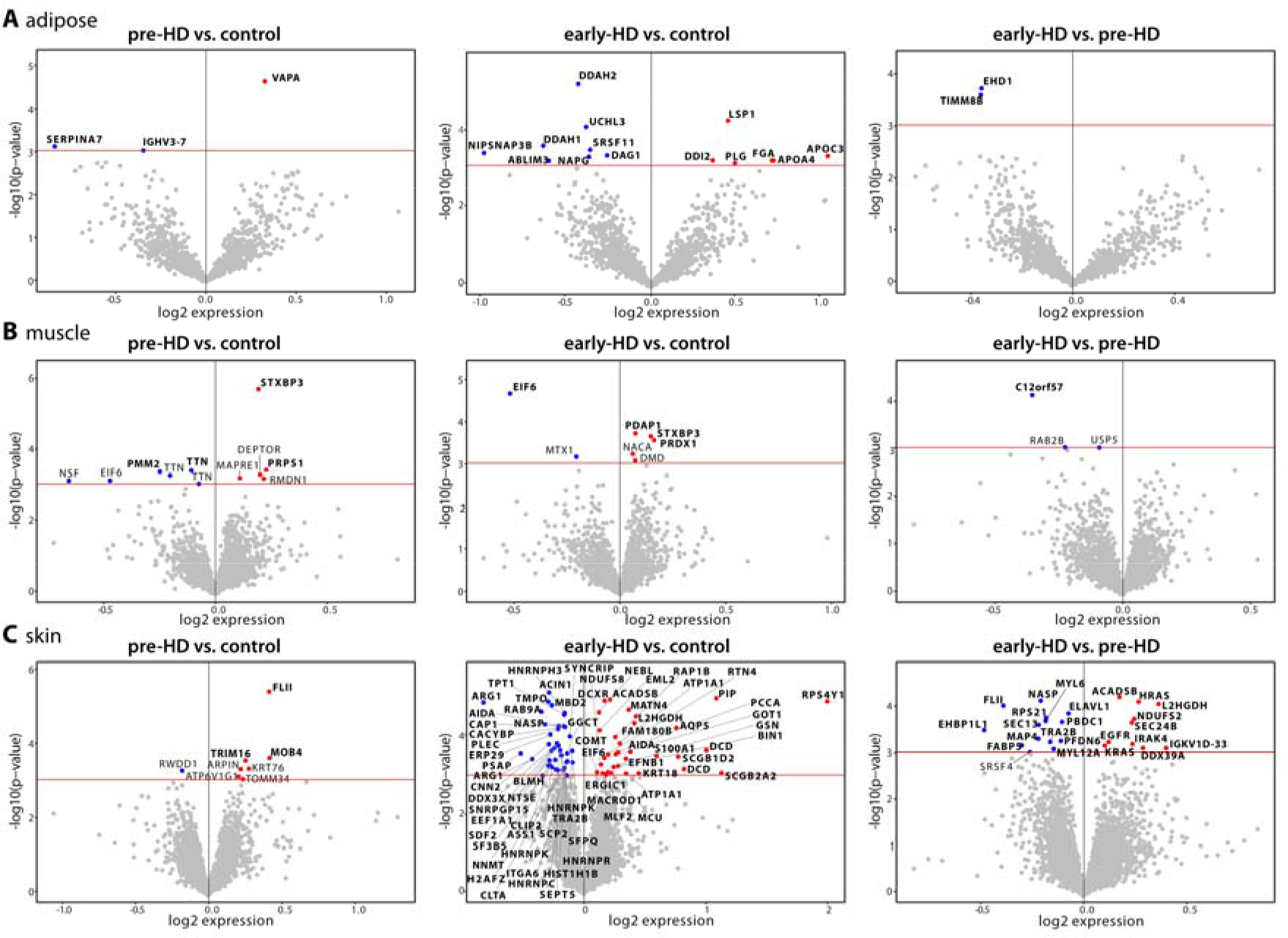
Proteomics analysis of adipose, muscle and skin samples. (**A-C**) Volcano plots of proteomics data from adipose (**A**), muscle (**B**) and skin (**C**) tissue samples. Pair-wise comparisons of pre-HD and early-HD against controls and early-HD against pre-HD are shown. The x-axis is the log_2_(expression) change and y-axis -lg_10_(*p*-value) from the ROTS analysis. The red horizontal line highlights the *p*-value of -lg_10_(3) = 0.001. Proteins with a significance value of less than 0.001 are depicted and proteins with an adjusted *p*-value (ROTS FDR) of < 0.05 are shown in bold. Up-regulated protein are shown in red, down-regulated proteins in blue. See also supplementary tables for full data.

In summary, for all three tissues we observed a progressive increase of the number of dysregulated proteins from pre-HD to early-HD stage. In adipose tissue there was no overlap of commonly dysregulated proteins in the three comparisons (Fig. 3A). In muscle, Syntaxin-binding protein 3 (STXBP3) was up-regulated in both, pre- and early-HD samples (Fig. 3B). We observed the largest changes in skin (Fig. 3C). Here, MOB family member 4, phocein (MOB4) was up-regulated in both, pre- and early-HD samples. 162 proteins were significantly (FDR < 0.05) dysregulated in the early-HD group. 23 proteins significantly changed their expression levels from pre-HD to early-HD stage (Fig. 3C).

Due to the low number of changed proteins, we only performed gene ontology enrichment in the adipose tissue dataset (early-HD compared to controls) and the skin dataset (early-HD compared to controls and the comparison of pre-HD to early-HD) (Table 3). Pathway and ontology analysis in the adipose tissue dataset hinted at dysregulated lipid metabolism and proteasome function (Table 3, adipose: early-HD vs. controls). Both have been extensively studied in HD. Peroxisome proliferator activated receptor alpha (PPARA) was strongly predicted as potential upstream regulator of these dysregulated proteins (Table 3). Worth noting, PPARA was also predicted as the most likely upstream regulator for the dysregulated gene analysis of the muscle dataset (Fig. 2B).

**Table 3.** Ontology Enrichment of proteomics data.

| Dataset | Pathways<br>1: WikiPathways 2019 (human)<br>2: KEGG 2019 (human) | Ontology<br>[biological process] | Regulators<br>1: ChEA 2016<br>2: TTRUST 2019<br>3: ENCODE 2015 |
| --- | --- | --- | --- |
| **Adipose:**<br>**early-HD**<br>**vs. control** | 1: Blood Clotting Cascade (1480.08), Statin Pathway (408.90), Proteasome Degradation (378.24)<br>2: Complement and coagulation cascades (546.42), Proteasome (225.26), Cholesterol metabolism (195.08) | chylomicron remodeling (2106.52), nitric oxide mediated signal transduction (1008.13), fibrinolysis (921.53), proteasomal ubiquitin-independent protein catabolic process (528.58), interleukin-1-mediated signaling pathway (209.32) | 1: HNF4A (14.55), AR (13.99), NANOG (13.59)<br>2: PPARA (273.66)<br>3: FOXA2 (29.83), NR2C2 (25.90), HNF4G (24.99) |
| **Skin:**<br>**early-HD**<br>**vs. control** | 1: Translation Factors (337.17), mRNA Processing (261.60), Methylation Pathways (234.97)<br>2: Spliceosome (298.11), Arginine biosynthesis (167.83), Tyrosine metabolism (72.16) | protein localization to cytoplasmic stress granule (356.98), mRNA splicing, via spliceosome (343.79), IRES-dependent viral translational initiation (198.37), aspartate family amino acid metabolic process (198.37), nucleoside metabolic process (198.37) | 1: XRN2 (23.62), AR (23.21), TP63 (22.90)<br>2: BRCA1 (34.50), VDR (21.17), AR (14.95)<br>3: NELFE (93.56), KAT2A (74.77), TAF7 (48.19) |
| **Skin:**<br>**early-HD**<br>**vs. pre-HD** | 1: Extracellular vesicle-mediated signaling in recipient cells (1346.70), ErbB Signaling Pathway (298.26), AGE/RAGE pathway (176.44)<br>2: ErbB signaling pathway (327.55), Oxytocin signaling pathway (295.74), Neurotrophin signaling pathway (205.61) | cargo loading into COPII-coated vesicle (1787.10), ERBB2 signaling pathway (942.71), ephrin receptor signaling pathway (130.95), Fc receptor signaling pathway (111.42), mRNA splicing, via spliceosome (65.83) | 1: SOX11 (22.22), EGR1 (22.10), FOXP2 (19.42)<br>2: TP53 (130.54), SP1 (9.04)<br>3: TEAD4 (60.02), GTF3C2 (23.76), SP2 (20.43) |

Ontology enrichment using the Enrichr website. See Methods for details. Only non-redundant, aggregated terms are shown. Only entries of human samples in the databases were considered. Only terms that applied to at least two dysregulated proteins were considered. The combined score is shown in brackets. Only entries with a combined score of 5 or greater were considered. Top three terms for pathways and regulators and top 5 terms for ontology are shown.

Strikingly, 21 out of the 28 dysregulated proteins in the adipose tissue dataset (early-HD vs. controls) are associated with the term ‘extracellular vesicle’. Extracellular vesicles (EVs) are composed of lipids, proteins and RNAs, are membrane coated and are secreted by most cells. Moreover, they are implicated in neurodegeneration, although the availability of data for HD is poor [24].

Pathway and ontology enrichment in the skin proteomics dataset showed that proteins related to gene expression in general are affected in the early-HD samples compared to controls (Table 3). RNA processing, translation, as well as amino acid metabolism seems to be dysregulated. The predicted regulators are all factors involved in gene expression. Of note, androgen receptor (AR) and lysine acetyltransferase 2A (KAT2A), the later in a complex with ataxin 7 (ATXN7), are both linked to polyglutamine diseases. A CAG expansion in AR causes spinal and bulbar muscular atrophy [25]; an expansion in ATXN7 causes spinocerebellar ataxia type 7 [26]. This could potentially indicate that there are common phenomena caused by a CAG repeat expansion, independent of the mutated gene. The enriched pathways in the comparison of the early-HD and pre-HD samples in the skin dataset (Table 3) are intertwined and regulate cell survival and proliferation. Consequently, TP53 (tumor protein P53), a key regulator of cellular survival, was predicted as an upstream regulatory protein. In addition, extracellular vesicle mediated signaling was again predicted with a huge combined score (Table 3).

## Discussion

Many valuable insights into the pathogenesis of HD have been gained from disease models, whether in mice, flies or non-human primates [27, 28] to suggest that the expression of the HD mutation affects the innate immune system and energy metabolism [29]. The in-depth molecular tissue signatures, which we derived from human HD patients’ peripheral tissues specifically indicate PPARA (peroxisome proliferator activated receptor alpha) as a regulatory protein with very high confidence for dysregulated genes in muscle in the pre-HD group (Fig. 2B) and for dysregulated proteins in adipose tissue in the early-HD group (Table 3). The family of PPA receptors are important regulators of various processes including inflammation, lipid and energy metabolism. The PPA receptors interact with e.g. PGC-1α (Peroxisome Proliferator-Activated Receptor Gamma Coactivator) to enable the correct function of both partners including in the central nervous system [30]. Paralogs of PPARA, namely PPARD [31, 32] and PPARG [33, 34] have been implicated in HD pathogenesis. Moreover, PPARs can be therapeutically targeted to improve phenotypes such as mitochondrial dysfunction in HD model systems [32-35].

The most consistently changed gene in the RNAseq data from muscle, adipose and fibroblasts was *TBC1D3D* (ENSG00000274419). In all tissues its expression was significantly down-regulated in the pre-HD group and restored towards control levels in the early-HD group (Fig. 2). TBC1D3D protein is expressed in all three tissue types (www.uniprot.org, [36]) and most probably acts as a GTPase activating protein to stimulate RAB5 function [37]. RAB5 is an early endosome marker and functions in endocytosis and membrane transport where it regulates receptor sorting and vesicle fusion [38]. Intriguingly, the HTT-HAP40 complex also acts as a regulatory protein complex for RAB5 [39]. Consequently, expression of mutant HTT leads to aberrant endosomal/lysosomal pathway function [40] and its secretion via a non-canonical pathway [41]. Endosomal pathway integrity is a prerequisite for functioning secretion of exosomes through multi-vesicular bodies (part of the endosomal system) and vice versa [42]. Exosomes are one of the three main species of extracellular vesicles (EVs), the others being microvesicles, also called ectosomes, and apoptotic bodies [43]. Apoptotic bodies form during cell death; microvesicles directly bud from the cell plasma membrane. All EVs are composed of lipids, proteins, RNAs and other small molecules. Their circulating nature makes them attractive messengers of information from the secreting cell towards the target cell. They are also able to cross the blood brain barrier and are found in virtually every bodily fluid [43]. When we analyzed the dysregulated proteins in the adipose dataset (early-HD vs. controls), we found an exceptionally high association of the gene ontology term ‘extracellular vesicle’ (for 21 out of the 28 dysregulated proteins) (Table 3). Additionally, gene ontology enrichment of dysregulated proteins in the skin dataset also pointed towards ‘Extracellular vesicle-mediated signaling in recipient cells’ (Table 3). Extracellular vesicles could play an important role in neurodegeneration as they have been implicated as one route for the spread of misfolded proteins, for instance in Parkinson’s disease, and they also contain other cargo such as miRNAs [44]. In HD, our human tissue data indicate that the expression of the HD mutation affects the homeostasis of extracellular vesicles similar to what has been observed before [45]. These findings certainly merit further investigation, in particular because EVs derived from peripheral tissues and biofluids are an attractive source of (peripheral) biomarkers for clinical trials.

Many different approaches aim to modify HD pathogenesis [46] with those targeting pathology close to the repeat expansion mutation probably having the best chances of success. Examples are those directly targeting the huntingtin RNA, thus reducing levels of the *HTT* mRNA and the encoded HTT protein (fragments) [8, 9, 47]. Indiscriminately lowering HTT levels, including the wild-type form, might lead to adverse consequences [8, 9]. For instance, *HTT* hypomorphic mutations, which result in lower expression of wild-type huntingtin, cause the developmental disorder Lopes-Maciel-Rodan syndrome [48]. Additionally, complete loss of wild-type HTT function led to behavioral deficits and progressive neuropathological changes in a mouse model of HD [49]. To circumvent these problems SNP-based HTT lowering approaches are underway to selectively target only the expanded allele harboring the SNP [8, 9, 47]. Our fully SNP phased fibroblasts and the iPSC lines derived from them can be used to analyze the feasibility of these approaches and to develop novel molecules for therapy. Our phylogenetic analysis of the SNP phasing data of our exclusively Caucasian HD patients indicate that some haplotypes might be less prone to expansions than others (Fig. 1C and D). While this data certainly would be more robust based on a larger sample size, similar observations have been reported previously [10, 12, 50, 51].

In summary, the biological signatures of the changes at RNA and protein levels point towards the involvement of inflammation, energy metabolism and vesicle biology in peripheral tissues in HD. It shows the potential to identify biological signatures from peripheral tissues in HD that could be suitable as biomarkers in clinical trials. In addition, we have generated a high-quality human reference data set and established valuable primary cell lines and iPSC cell lines from participants taking part in a large longitudinal observational study, Enroll-HD, with deep phenotyping [52]. These provide an important resource for further research into, for instance, somatic repeat instability, cell type specific effects of mutant HTT or the HD epigenome [5]. iPSC lines are also valuable tools in many neurodegenerative disorders [53], including HD [54], and our set of well-characterized lines can be used to study pathomechanisms across diseases.

## Materials and Methods

### Study participants

Participants of the MTM-HD study were recruited at the departments of neurology of Ulm University (Germany) and University College London (United Kingdom). At both institutions the study followed a standardized protocol. Participants were only included if there were no contra-indications to muscle biopsy, e.g. a clotting disorder or evidence to suggest cardio-respiratory abnormalities. For HD patients, the CAG-tract length was determined and participants were clinically assessed as described for the TrackOn and TRACK-HD studies [55, 56]. Clinical assessment included the United Huntington Disease Rating Scale (UHDRS) motor part to derive the total motor score and the UHDRS total functional capacity scale (TFC) [57]. The disease burden score (DBS) was calculated from each HD participant’s CAG repeat length and age according to the following formula: (CAG-35.5) x age [58]. A DBS of 250 or greater was used to screen for potential HD participants. Furthermore, HD participants were categorized as pre-HD if they had a diagnostic confidence level score of 2 or less on the UHDRS motor scale, or as early-HD if they were in TFC stages of 1 or 2, indicative of early motor manifest HD. Only participants with no signs of neuromuscular disease on clinical examination were enrolled. The local ethics committees at Ulm University and University College London approved the study (Ulm: 265-12; London: 12/LO/1565), and written informed consent was obtained from each participant. Following informed consent, participants were given a 9-digit unique identifier as described [59].

### Tissue biopsy and sample collection

In order to ensure samples are collected and processed in exactly the same manner in Ulm and in London, personnel at either site received on-site training on all aspects of the protocol. All samples were collected following an overnight fast (water was permitted) between 7.30-9 am local time (CET or GMT) to account for any influence of circadian rhythm. Open biopsies of the skin, subcutaneous adipose tissue and *M. vastus lateralis* were obtained following local anesthesia. Participants were contacted on the day after and 7 days following the procedures. After one week, the stitches were removed. Tissues were dissected (skeletal muscle, adipose tissue and skin), immediately snap-frozen in liquid nitrogen and stored at -80°C. Blood samples were taken following informed consent on the screening day. This included routine blood tests (full blood count, renal, liver, bone and clotting profiles, creatinine kinase and group and save). In addition, plasma and buffy coat were separated. PBMCs were extracted from CPT/heparin tubes (BD Vacutainer CPT) as previously described [60]. Cell pellets were then snap frozen in liquid nitrogen and stored at -80°C. For histochemistry and immunohistochemistry the muscle sample was mounted on a piece of cork in TissueTek with fibers oriented perpendicular to the cork and then snap-frozen in liquid nitrogen-cooled 2-methylbutane and stored at -80°C. For electron microscopic analysis a muscle fiber was pinned onto a cork plate with two needles in the operating theatre and immediately fixed in 2.5% glutaraldehyde.

### Generation of primary cell lines

Primary human fibroblast and myoblast cell lines from the MTM participants were established and maintained as previously described [61].

### RNA and DNA extraction

RNA extraction from tissues was performed by Labcorp (NC, US) using the Qiagen miRNeasy kit. RNA had to pass the following specifications from an Agilent Bioanalyzer run to be used for subsequent sequencing: 28S/18S ratio: 0.8 – 3.0 and RIN Score: ≥ 6.0. For fibroblast lines, RNA and DNA were extracted simultaneously by Q Squared Solutions Expression Analysis LLC (NC, US). The Qiagen AllPrep DNA/RNA kit was used to simultaneously purify genomic DNA and total RNA from the same sample. Briefly, samples were lysed in a guanidine-isothiocynate containing buffer and the lysate was passed through an AllPrep DNA spin column which selectively binds genomic DNA. The column was washed and the genomic DNA was eluted in EB buffer stored at -20°C. Ethanol was added to the flow-through of the AllPrep DNA spin column which allows for efficient binding of RNA. The mixture was then applied to a RNeasy spin column (Qiagen), where total RNA binds to the membrane. RNA was then eluted in RNase-free water, quantified and integrity assessed using the RNA 6000 Nano Assay on a Bioanalyzer 2100. RNA passed QC with RIN values ≥ 7.0. The purified RNA was stored at - 80°C.

### Methylation analysis of fibroblast derived DNA

Methylation analysis was performed by Q Squared Solutions Expression Analysis LLC (NC, US). DNA samples were assayed for cytosine methylation profiles using the Infinium Methylation Assay, essentially as described by the manufacturer. 500 ng of input DNA was treated with sodium bisulphite to convert unmethylated cytosines to uracil, leaving methylated cytosines unchanged. The treated DNA sample was then denatured, neutralized, and isothermally amplified. The amplified DNA was fragmented, precipitated with isopropanol and re-suspended prior to hybridization onto BeadChips. The converted and non-converted amplified DNAs hybridize to their corresponding probes, and excess DNA was washed away. Hybridized DNAs undergo single-base extension and staining for labelling, followed by scanning on an Illumina iScan instrument for detection.

### RNA sequencing of adipose and muscle tissue RNA

RNA sequencing was performed by Q Squared Solutions Expression Analysis LLC (NC, US). RNA samples were converted into cDNA libraries using the Illumina TruSeq Stranded Total RNA sample preparation kit (Illumina # RS-122-2303). Briefly, total RNA samples were concentration normalized, and ribosomal RNA (rRNA) was removed using biotinylated probes that selectively bind rRNA species. The resulting rRNA-depleted RNA was fragmented using heat in the presence of divalent cations. Fragmented RNA was converted into double-stranded cDNA, with dUTP utilized in place of dTTP in the second strand master mix. A single ‘A’ base was added to the cDNA and forked adaptors that include index, or barcode sequences were attached via ligation. The resulting molecules were amplified via polymerase chain reaction (PCR). During PCR the polymerase stalls when a dUTP base is encountered in the template. Since only the second strand includes the dUTP base, this renders the first strand the only viable template, thereby preserving the strand information. Final libraries were quantified, normalized and pooled. Pooled libraries were bound to the surface of a flow cell and each bound template molecule was clonally amplified up to 1000-fold to create individual clusters. Final libraries were sequenced on the HiSeq Illumina sequencing platform with paired-end, 50 bp long reads with a total read depth of approximately 150M reads per sample. Sequencing raw files were demultiplexed, adapter trimmed and clipped for low quality base calls. Final fastQ files were produced with reads of a minimum length after clipping/trimming of 25 nucleotides.

### RNA sequencing of fibroblast RNA

RNA sequencing was performed by Q Squared Solutions Expression Analysis LLC (NC, US). Sequencing libraries were created using the Illumina TruSeq Stranded mRNA method (Illumina, RS-122-2103), which preferentially selects for messenger RNA by taking advantage of the polyadenylated tail. In summary, approximately 100 ng of total RNA per sample was used to purify poly-adenylated RNAs using oligo-dT attached to magnetic beads. Purified mRNAs were fragmented using heat in the presence of divalent cations. The fragmented RNAs were converted into double-stranded cDNA, with dUTP utilized in place of dTTP in the second strand master mix. This facilitates the preservation of strand information, as amplification in the final PCR step will stall when it encounters Uracil in the nucleotide strand, rendering the first strand as the only viable amplification template. The double stranded cDNA underwent end-repair, A-tailing, and ligation of adapters that include index sequences. The final libraries were amplified via polymerase chain reaction (PCR), after which they were quantified, normalized and pooled in preparation for sequencing. Normalized libraries were multiplexed for efficient sequencing to the required number of reads per sample. Pooled libraries were bound to the surface of a flow cell, and each bound template molecule was clonally amplified up to 1000-fold to create individual clusters. Libraries were sequenced using the Illumina sequencing-by-synthesis platform, with a sequencing protocol of 100 bp paired-end sequencing and total read depth of 40M reads per sample on a NovaSeq. Sequencing raw files were demultiplexed, adapter trimmed and clipped for low quality base calls. Final fastQ files were produced with reads of a minimum length after clipping/trimming of 25 nucleotides.

### Bioinformatics analysis of RNAseq datasets

FastQ files were quality checked with FastQC v0.11.5 [62]. All files passed QC. The reads were aligned against Ensembl *homo sapiens* GRCh38 release 90 using STAR aligner v2.5.3a [63]. Reads were quantified using salmon v0.8.2 [64]. Samples were screened for outliers using a combined PCA and clustering analysis. A sample was defined as an outlier if it was outside a 68% probability ellipse in PCA analysis and was outside a -2.5 standardized connectivity cutoff of an Euclidean distance matrix of all samples per each tissue [65]. This procedure identified 6 samples in the adipose (n = 2 for each control, pre-HD and early-HD), 3 samples in the muscle (n = 1 control, n = 2 pre-HD) and 10 samples in the fibroblast (n = 4 control, n = 4 pre-HD, n = 2 early-HD) datasets. For all subsequent analyses, both RNAseq batches for muscle and adipose tissues were combined. Transcriptional dysregulation was computed using tximport v1.10.0 [66] and DESeq2 v1.22.1 [67]. The DESeq2 intercept design normalized counts are available as supplementary information files. For the dysregulation analysis HD clinical stage (control, pre-HD, early-HD) was used as the variable of interest. RNAseq batch (for tissues), gender, site of sampling, age and BMI were used as covariates in the modeling. We factorized the two continuous variables (age and BMI) into 5 intervals before evaluation with the cut function in R. Ashr was used as the fold change shrinkage estimator [68]. DESeq2 analysis files are available as supplementary information files. Ontology analysis was carried out using the Enrichr website [69, 70].

### Proteomic analysis and bioinformatics

Proteomics analysis was performed by proteome sciences (www.proteomics.com). For TMT labelling (Thermo Fisher), 2 mg of muscle tissue were reduced (dithiothreitol), alkylated (iodoacetamide), digested (trypsin) to generate peptides, desalted (SepPak tC18 cartridges (Waters, Milford, MA, USA)) and lyophilized. Peptides were mixed with their respective TMT 10plex tag and incubated for 1 hour at room temperature. Individual TMT reactions were terminated with hydroxylamine and the labelled digests of each of the 10 samples were pooled into the respective TMT 10plex and incubated for another hour. Finally, the TMT 10plex analytical sample was acidified, diluted to an acetonitrile concentration less than 5%, divided into two aliquots, desalted and lyophilized to completion. Each TMT 10plex analytical sample aliquot was separated into 12 × 4-minute fractions by strong cation exchange chromatography (polySULFOETHYL-A column (PolyLC) and HPLC system (Waters Alliance 2695)) and the 12 fractions combined by smart pooling into six fractions with roughly equal peptide amounts for subsequent processing. The muscle samples were subjected to two SCX-runs as the total load of the muscle TMT 10plexes was 20 mg each and makes two runs necessary. Re-suspended peptides were loaded onto a nanoViper C18 Acclaim PepMap 100 pre-column (Thermo Scientific). Each fraction was analyzed in duplicate by LC-MS/MS using the EASY-nLC 1000 system coupled to an Orbitrap FusionTM TribridTM Mass Spectrometer (both Thermo Scientific) for each of the TMT 10plex sets. Peptide mass spectra were acquired throughout the entire chromatographic run (180 minutes). MS-runs across the TMT 10plexes were submitted to Proteome Discoverer (PD) v1.4 (Thermo Scientific) using the Spectrum Files node. Spectrum selector was set to its default values while the SEQUEST HT node, was suitably set up to search data against the human FASTA UniProtKB/Swiss-Prot database (version 9606_Human_01082016 (created August 2016)). The reporter ions quantifier node was set up to measure the raw intensity values for TMT 10plex mono-isotopic ions (126, 127N, 127C, 128N, 128C, 129N, 129C, 130N, 130C, 131). The SEQUEST HT search engine was programmed to search for tryptic peptides (with up to two missed cleavages allowed) and with static modifications of carbamidomethyl (C), TMT6plex (K), and TMT6plex (N-Term). Dynamic modifications were set to deamidation (N/Q), oxidation (M), and phosphorylation (STY). Precursor mass tolerance was set to 20 ppm and fragment (b and y ions) mass tolerance to 0.02Da. The first stage of data processing was performed using the SQuaT Bioinformatics Module 3.1.1. Each TMT 10plex set was filtered to include PSMs (peptide spectral matches) that had a signal in at least one out of 10 TMT reporter ion channels. A correction procedure was then implemented to address isotopic impurities. Isotope correction factors used in this procedure were specific to the TMT batch used for labelling. Next, for every mass spectrometry run, the TMT reporter ion intensities were normalized using the sum-scaling technique at the PSM level. Briefly, for each TMT channel all reporter ion intensities were summed and the median value across all summed channels was calculated. A correction factor was obtained by dividing the individual summed reporter ion intensity for a channel by the median of all channels. Finally, each individual PSM ion intensity was multiplied by the correction factor for the relevant TMT reporter channel. Post sum-scaling, the ratios of each sample PSM was related to the reference sample channel. The PSM level ratios to the reference sample were calculated as a log2 transformation of sample PSM intensity relative to the reference sample PSM intensity. The data quality of the samples was controlled using quality control charts. Two quality control metrics per sample were calculated: the median (measure of central tendency) and the inter-quartile range (IQR) (measure of scale) using peptide and protein distributions. Both metrices are robust and stable to outliers and are therefore are good for detecting extreme outliers. The median log expression and inter-quartile range (IQR), were traced using control charts for all datasets on peptide and protein level. A sample was considered as a strong outlier if either QC metric value was more than three standard deviations from the overall mean. A second stage of data processing was performed using the FeaST Bioinformatics Module 1.3.1. Imputation was performed to obtain quantitative values for the remaining missing data points using an iterative PCA method [71]. Following imputation, the data were normalized across the sets using quantile normalization method to center the sample values across the different TMT 10plex sets. Multifactorial limma [72] modelling was used to remove TMT 10plex set and site of sampling as batch effects. The final datafiles for adipose, muscle and skin proteomics analysis are available as supplementary information files.

### Bioinformatics analysis of proteomics datasets

For further analysis of the peptide datasets, all non-unique entries for peptides (shared_status_Gene) were removed, followed by removal of rows with unassigned protein IDs. We then used ‘Peptide’ as the identifier to collapse rows (WGCNA function [73]) onto unique peptides entries. We next employed the same strategy as described above to identify outliers in the peptide and protein datasets, but could not detect any outlying samples. Following this, we consecutively removed batch effects with limma [72] for both, peptide and aggregated protein datasets. Adipose: Gender, BMI, age; Muscle: BMI, age; Skin: Age, gender, BMI. To compute peptide and protein dysregulation, we used ROTS [23]. For each pairwise comparison we used the respective full dataset for modeling and ran 1000 bootstraps. Reproducibility Z-score value was greater than 2 for all comparisons. The ROTS analysis files are available as supplementary information files.

### SNP sequencing

Genomic DNA extraction, sample QC, library preparation, sequencing reactions, and bioinformatics analyses were conducted at GENEWIZ, Inc. (South Plainfield, NJ, USA). High molecular weight genomic DNA was extracted from 2-4 million cells per sample, using Qiagen Genomic-tip 100/G HMW Kit (Qiagen, Hilden, Germany), according to manufacturer’s protocol. Sample amount was quantified using a Qubit 2.0 Fluorometer (Invitrogen, Carlsbad, CA, USA), and sample purity and integrity was checked using a Nanodrop spectrophotometer and a pulsed field gel analysis (PFGE), or an Agilent TapeStation (Palo Alto, CA, USA), respectively. All samples were processed through the Chromium Controller following the standard manufacturer’s specifications. DNA phasing libraries were generated from 1.07-2.41 ng of DNA per sample, using the 10X Genomics Microfluidic Genome Chip and 10X Genomics Chromium Genome kit (10X Genomics, CA, USA), according to manufacturer’s protocol. The sequencing libraries were evaluated for quality using an Agilent TapeStation (Palo Alto, CA, USA), and quantified by using a Qubit 2.0 Fluorometer (Invitrogen, Carlsbad, CA, USA), and qPCR (Applied Biosystems, Carlsbad, CA, USA). The pooled sequencing libraries were loaded onto an Illumina HiSeq 4000 sequencer to achieve 150 Gb of data per sample. The samples were sequenced in a configuration compatible with the recommended guidelines as outlined by 10X Genomics (2 × 150 bp configuration, with 8 bp single index). A single sample with insufficient phasing results was sequenced to an additional 300 Gb of data. Raw sequence data (bcl) files generated by the machine were converted into fastq files and de-multiplexed using the 10X Genomics Cell Ranger software. Genome sequence variant calling, phasing and structural variant calling was done using 10X Genomics Long Ranger software with interactive visualization on the Loupe genome browser.

### HD haplotyping

The definitions of the most common 16 haplotypes in HD subjects were described previously [10]. The here described HTT haplotypes were constructed using 20 SNPs and 1 insertion-deletion polymorphism (rs149109767), and were named based on the frequency in the HD subjects with European ancestry. The 16 most frequent haplotypes account for approximately 90% and 82% of disease and normal chromosomes in HD subjects with European ancestry, respectively. We defined HTT haplotypes further based on the 1000 Genomes Project (KGP) data (https://www.internationalgenome.org/; phase 3 data). Briefly, we extracted the same haplotype-defining 21 variations from all of the phased KGP data (5008 chromosomes), and constructed haplotypes subsequently. We then calculated the frequency in KGP data (all populations) for unique haplotype. Excluding the 16 haplotypes, which we had defined previously [10], we sorted haplotypes based on frequency and assigned names accordingly. Finally, 253 additional haplotypes were defined. The full list of the newly defined HD haplotypes is available in the supplementary information. HTT haplotypes were clustered via binary distance. A dendrogram was generated using Ward’s linkage. This analysis was performed separately for the normal and mutant haplotypes. In each dendrogram, three clusters were identified via k-means clustering represented by different colors. Analysis was performed in R using packages circlize [74] and dendextend [75]. Barplots representing haplotype frequencies were generated via ggplot2 [76].

### Data, code and script availability

All analyses were conducted in R v3.5.2, v3.6.1 and v4.0.3 [77]. Further details about the bioinformatics evaluation, as well as scripts and code are available upon request from A.N. The datasets supporting the conclusions of this article are included within the article and its additional files.

## Supporting information

MTM_all_metadata

MTM-HD_HD_haplotype_definitions

MTM-HD_HTT_SNP_haplotyping_fibroblasts

MTM_proteomics_adipose_peptide

MTM_proteomics_adipose_peptide_ROTS_analysis

MTM_proteomics_adipose_protein

MTM_proteomics_adipose_protein_ROTS_analysis

MTM_proteomics_muscle_peptide

MTM_proteomics_muscle_peptide_ROTS_analysis

MTM_proteomics_muscle_protein

MTM_proteomics_muscle_protein_ROTS_analysis

MTM_proteomics_skin_peptide

MTM_proteomics_skin_peptide_ROTS_analysis

MTM_proteomics_skin_protein

MTM_proteomics_skin_protein_ROTS_analysis

MTM_RNAseq_adipose_outliers_removed_DESeq2_analysis

MTM_RNAseq_adipose_outliers_removed_normalized_counts

MTM_RNAseq_fibroblasts_outliers_removed_DESeq2_analysis

MTM_RNAseq_fibroblasts_outliers_removed_normalized_counts

MTM_RNAseq_muscle_outliers_removed_DESeq2_analysis

MTM_RNAseq_muscle_outliers_removed_normalized_counts

## Author Contributions

Conceptualization: S.J.T. and M.O.; Methodology: A.N., M.O.; Software: A.N.; Validation: A.N., K.K., T.H., J.P., S.J.T., M.O.; Formal Analysis: A.N; Investigation: A.N., K.K., T.H., N.B., J.H., S.T., J.P., H.S., S.H., S.J.T., M.O.; Resources: A.N., D.J.L., J.M.L., S.J.T., M.O.; Data Curation: A.N., M.F., M.O.; Writing – Original Draft: A.N., M.O.; Writing – Review & Editing: All authors; Visualization: A.N.; Supervision: S.J.T., M.O.; Project Administration: S.J.T., M.O.

## Acknowledgments

First and foremost, we thank all participants of the study for their time and their willingness to undergo tissue biopsies. The authors acknowledge support by the High Performance and Cloud Computing Group at the Zentrum für Datenverarbeitung of the University of Tübingen, the state of Baden-Württemberg through bwHPC and the German Research Foundation (DFG) through grant no INST 37/935-1 FUGG.

## Supplemental material

**MTM_all_metadata.xlsx:** All identifiers and demographic and clinical data

**RNAseq datafiles:** Normalized count files for adipose, muscle and fibroblasts

- MTM_RNAseq_adipose_outliers_removed_normalized_counts.xlsx
- MTM_RNAseq_muscle_outliers_removed_normalized_counts.xlsx
- MTM_RNAseq_fibroblasts_outliers_removed_normalized_counts.xlsx

**RNAseq evaluation files:** DESeq2 analysis files including fold change and *p*-values

- MTM_RNAseq_adipose_outliers_removed_DESeq2_analysis.xlsx
- MTM_RNAseq_muscle_outliers_removed_DESeq2_analysis.xlsx
- MTM_RNAseq_fibroblasts_outliers_removed_DESeq2_analysis.xlsx

**Proteomics datafiles**: Peptide and protein expression files for adipose, muscle and skin

- MTM_proteomics_adipose_peptide.xlsx
- MTM_proteomics_adipose_protein.xlsx
- MTM_proteomics_muscle_peptide.xlsx
- MTM_proteomics_muscle_protein.xlsx
- MTM_proteomics_skin_peptide.xlsx
- MTM_proteomics_skin_protein.xlsx

**Proteomics evaluation files**: ROTS analysis files including fold change and *p*-values

- MTM_proteomics_adipose_peptide_ROTS_analysis.xlsx
- MTM_proteomics_adipose_protein_ROTS_analysis.xlsx
- MTM_proteomics_muscle_peptide_ROTS_analysis.xlsx
- MTM_proteomics_muscle_protein_ROTS_analysis.xlsx
- MTM_proteomics_skin_peptide_ROTS_analysis.xlsx
- MTM_proteomics_skin_protein_ROTS_analysis.xlsx

***HTT* haplotyping:** Full list of HD haplotypes and *HTT* alleles SNP information

- MTM-HD_HD_haplotype_definitions.xlsx
- MTM-HD_HTT_SNP_haplotyping_fibroblasts.xlsx

